# Mapping the genomic landscape of peach and almond with PrunusMap

**DOI:** 10.1101/2024.03.26.586732

**Authors:** Najla Ksouri, María Ángeles Moreno, Bruno Contreras-Moreira, Yolanda Gogorcena

## Abstract

High throughput sequencing is accelerating plant breeding. Genotyping, quantitative trait locus or genome-wide association analyses are essential to locate alleles of interest in germplasm collections. These applications require accurately mapping nucleotide markers within growing collections of reference genome sequences. PrunusMap (https://prunusmap.eead.csic.es) is a user-friendly Web application designed to facilitate these tasks to *Prunus* breeders. It efficiently aligns markers and locates SNPs on peach and almond genomes, providing lists of nearby genes and curated proteins. It can align all markers from a genetic map in one step, producing their physical positions. By searching multiple maps, it can also solve the problem of finding equivalent gene identifiers across different annotations. Three intuitive tools, “Find markers”, “Align sequences” and “Locate by position”, are available to accelerate mapping analyses, particularly for users with limited bioinformatics expertise. New genomes, annotations or marker sets will be added based on their interest to the community.

## 1. Introduction

In the Rosaceae family, the availability of over 62 whole genomes and annotations provide robust foundation for marker development (https://www.plabipd.de/plant_genomes_pa.ep, last accessed November 2023). Restriction Fragment Length Polymorphism (RFLP) markers were instrumental in producing a saturated map for almond, the first and most comprehensive in the stone fruit genus *Prunus* (Viruel et al., 1995). Subsequently, randomly amplified polymorphic DNA (RAPD) markers found widespread use in germplasm diversity studies in peach and other *Prunus* species, helping map loci controlling traits such as flesh color and fruit texture (Araújo et al., 2010). The development of amplified restriction fragment length polymorphism (AFLP) markers revealed associations with traits such as resistance root-knot nematodes (Gillen & Bliss, 2005) and evergreen (*evg*) (Wang et al., 2002). However, the low reproducibility (RAPDs) and high costs (RFLPs, AFLPs) of these markers led to their replacement by SSRs and SNPs. While SSR markers find frequent utility in *Prunus* breeding, SNP markers have gained prominence due to their cost-effectiveness, high-throughput and genome-wide coverage (Aranzana et al., 2019; Butiuc-Keul et al., 2022). Moreover, the accurate prediction of marker positions and the identification of nearby genes are critical for understanding the genetic mechanisms underlying target traits and accelerating modern breeding cycles. For instance, research on Sharka disease in peach identified three highly significant associated SNPs on chromosome 2 and 3, conferring reduction in susceptibility to Plum pox virus. The *Prupe*.*2G065600* gene on chromosome 2, encoding an RTM2-like was selected as a major effect candidate gene (Cirilli et al., 2017). The emergence of SNP markers has indeed revolutionized *Prunus* breeding with two main repositories cataloging the genetic variants: the PeachVar-DB portal (Cirilli et al., 2018) and the Rosaceae Genome Database (GDR) (Jung et al., 2004, 2008, 2019). The PeachVar-DB portal provides different tools to retrieve SNPs and Indels extracted from whole genome sequences libraries of peach and wild relatives (Cirilli et al., 2018). Users can conveniently access these variants by specifying either a specific gene identifier or a genomic region of interest, with all coordinates extracted from the peach reference genome version 2.0. In contrast, the GDR stands out as a more multifaceted database, encompassing a broad spectrum of genomic and genetic data within the Rosaceae family. It provides a diverse range of tools aimed at exploring these resources. However, it lacks in capturing high-density genomic data obtained from re-sequencing projects (Cirilli et al., 2018). Moreover, browsing the GDR often requires switching between multiple pages, which can be cumbersome.

A friendly tool to map the location of genetic markers rapidly and accurately, along with information about nearby genes, could assist breeders, particularly those with limited bioinformatics expertise. To address this need, we created PrunusMap. Its objective is to streamline the process of locating *Prunus* markers on both genetic and physical maps, accommodating various input formats. While initially focused on *Prunus persica* and *Prunus dulcis*, it can be extended to other *Prunus* species. The Web application is accessible at https://prunusmap.eead.csic.es and offers three features to retrieve data:

1. “Find markers”: to retrieve the position of markers by providing their identifiers.
2. “Align sequences”: to obtain the position of FASTA sequences by pairwise alignment.
3. “Locate by position”: to examine specific loci by map position.

It is a fork of a pre-existing Webtool called Barleymap, originally designed to serve the barley community (Cantalapiedra et al., 2015), confirming that it can be tailored to any plant species with published genetic and genomic maps.

## 2. Material and methods

### 2.1 PrunusMap Web-interface

PrunusMap is a freely accessible application created as a fork of Barleymap (Cantalapiedra et al., 2015). Its back-end functionality and interactivity are implemented in Python 2.6, relying on CherryPy to handle the user requests (Hellegouarch, 2007). The front-end interface uses a Perl graphical library for chromosome visualization and intuitive interaction (https://github.com/pseudogene/genetic-mapper). The data flow and architecture are summarized in **Figure 1**.

**Figure 1.**
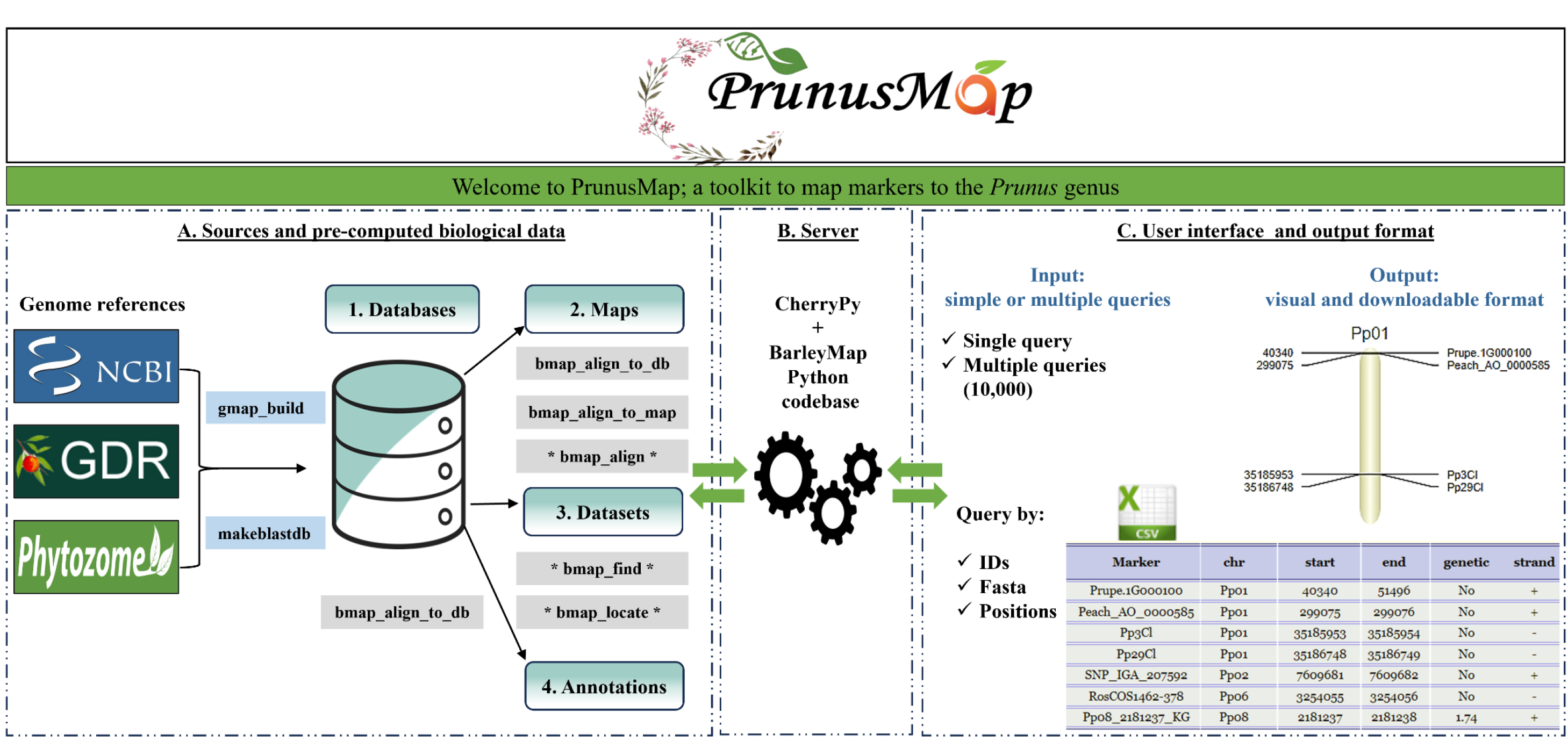
Workflow of PrunusMap Web application. The reference genome sequences of *Prunus persica* and *Prunus dulcis* were downloaded from NCBI, GDR and Phytozome. Gmap_build and makeblastdb command line executables were used to build the corresponding databases from the FASTA files. Databases, maps, datasets, and annotations represent the essential types of biological data required for PrunusMap configuration. Grey-highlighted boxes correspond to tools that are activated once each of the biological resources is properly set up (exp: “bmap_align_to_db” tool is activated once the database is well configured). Those tagged with asterisks correspond to Web-accessible tools, while the rest correspond to the standalone commands. PrunusMap accepts both simple or multiple queries as input and the results are displayed in visual and downloadable format (CSV).

### 2.2 Biological resources

PrunusMap stores and categorizes data into four different classes: databases, maps, datasets and annotations.

#### 2.2.1 Databases

A database is a genome sequence in FASTA format supporting the sequence alignments. Peach and almond references were sourced from NCBI (Sayers et al., 2022), Phytozome (JGI) (Goodstein et al., 2012) and GDR (Genome Database for Rosaceae) (Jung et al., 2004, 2008, 2019) then indexed from GMAP and BLASTN using “gmap_build” and “makeblastdb” (Boratyn et al., 2013; Wu & Watanabe, 2005). An example of database configuration is provided in **Figure S1**.

Currently, PrunusMap hosts several available reference genomes, described in **Figure 2**.

**Figure 2.**
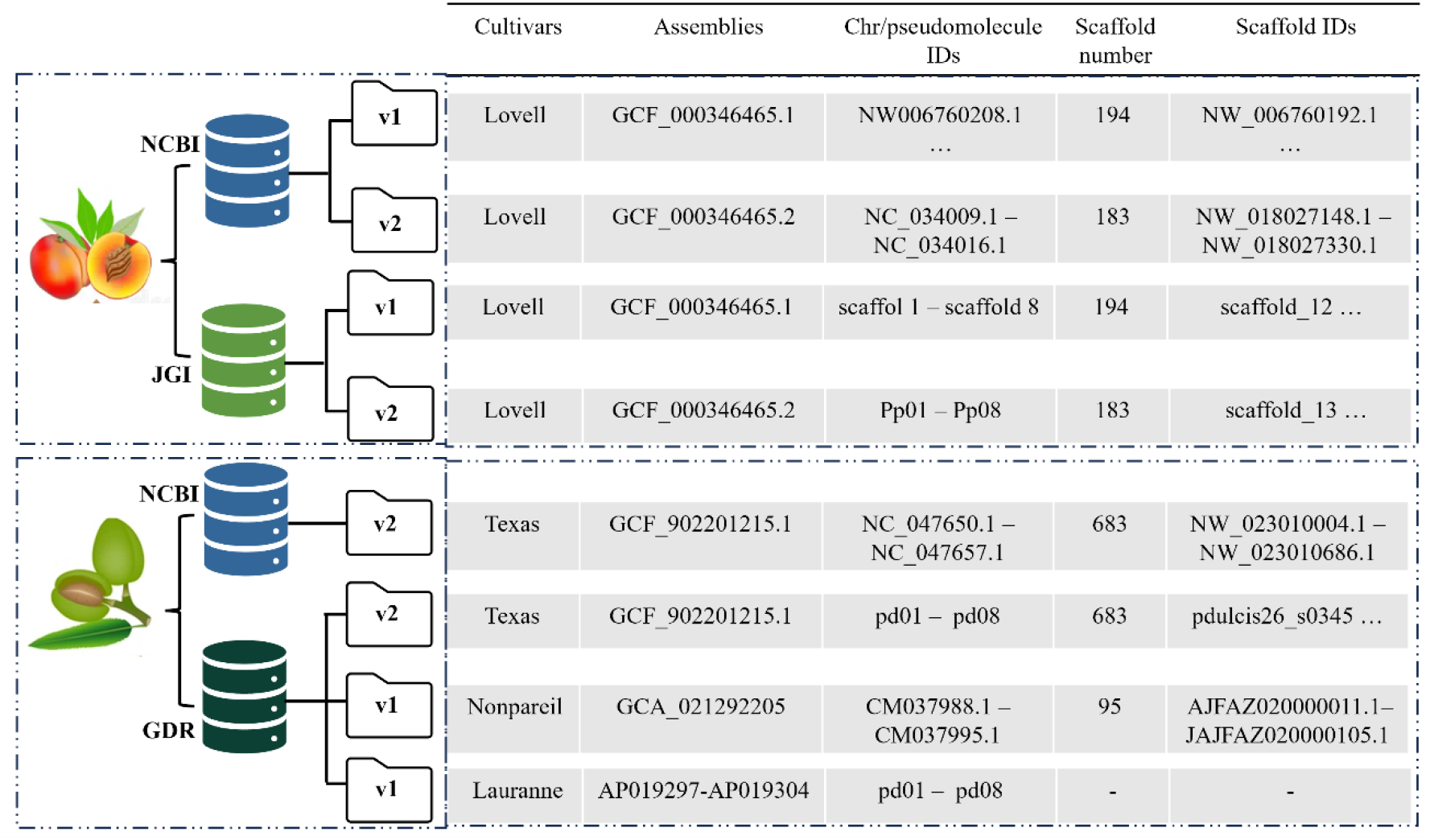
Illustration of PrunusMap databases. Please note that when “…” is used means that the IDs are not in continued order. However, when a range is used; for instance, NC_034009.1 – NC_034016.1 means the IDs are named in consecutive order. Chr refers to chromosomes.

#### 2.2.2 Maps

A map is a file type designed to store the positional arrangement, whether physical or genetic, of sequences derived from databases. PrunusMap includes eight maps:

- Pp_NCBI_V1, Pp_NCBI_V2, Pp_JGI_V1, Pp_JGI_V2, are physical maps associated with the distinct *Prunus persica* databases.
- Pd_Texas_NCBI_V2, Pd_Texas_GDR_V2, Pd_Nonpareil_GDR_V1 and Pd_Lauranne_GDR_V1 correspond to physical maps associated with the distinct *Prunus dulcis* databases.

#### 2.2.3 Datasets

PrunusMap datasets are genes, molecular markers and UniProt proteins, often associated with AlphaFold structural models (Li, 2023). Each dataset is a collection of one of these classes along with their precomputed map positions, determined through sequence alignment against the reference database. Genes and markers were aligned using GMAP and/or BLASTN while proteins were mapped with miniprot (Li, 2023). Datasets are crucial in identifying and tracking genetic markers across different peach and almond cultivars. They also facilitate the exploration of neighboring loci. See **Table 1** for a summary of gene models, genetic markers and lifted-over proteins.

**Table 1.**
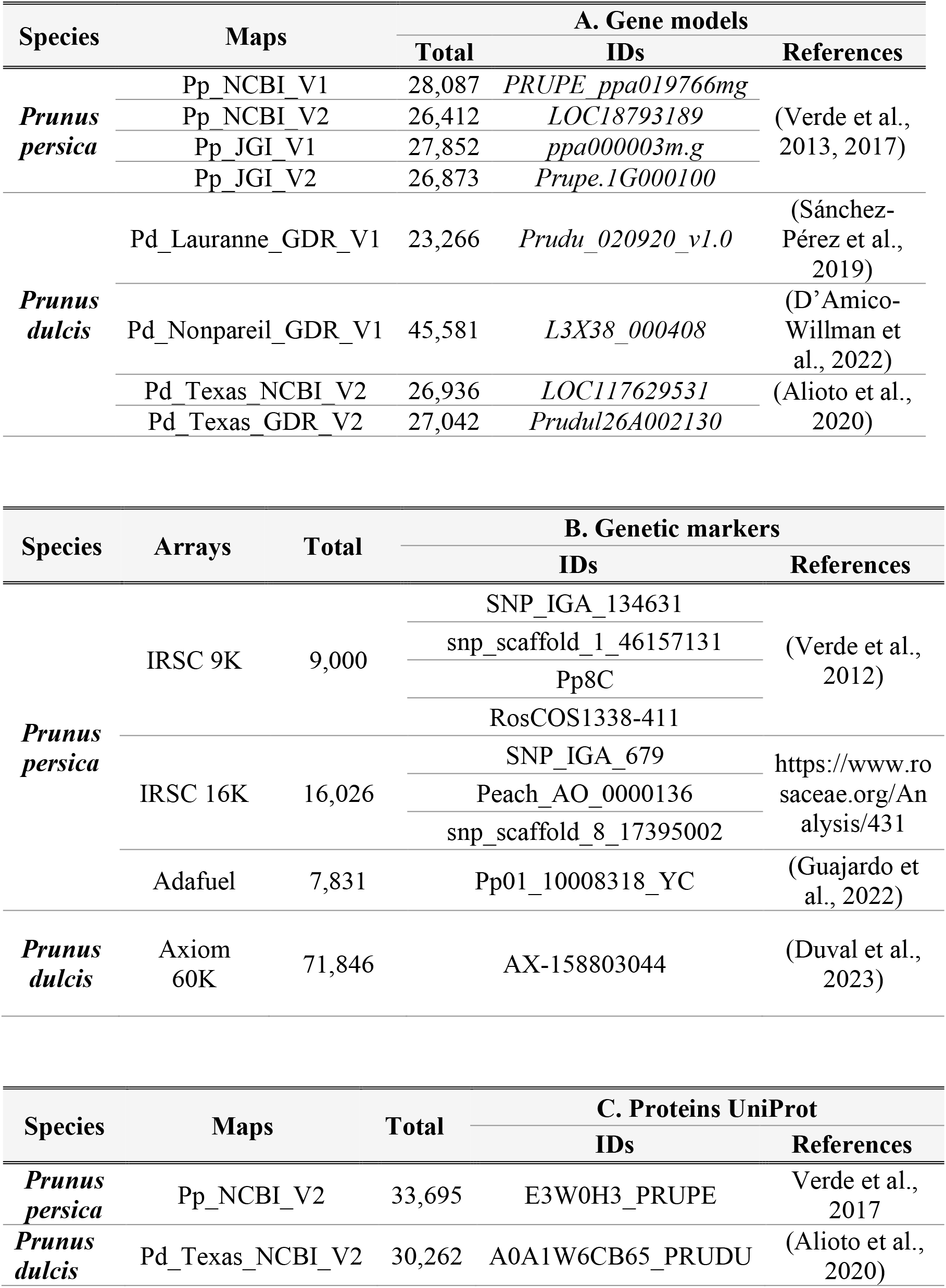
Gene models **(A)** markers **(B)** and mapped protein sequences **(C)** in peach and almond cultivars

The IRSC 9K and IRSC 16K SNP arrays for peach were developed by the International Rosaceae SNP Consortium (IRSC) and were downloaded from the GDR (https://www.rosaceae.org/organism/24333). The 9K array was obtained after resequencing 56 peach accessions and the 16K chip built upon it (Gasic et al. 2022).

Genetic markers for the hybrid peach-almond rootstock “Adafuel” were retrieved from the maps resulting of the analysis of the “Adafuel × Flordaguard” population (Guajardo et al., 2022). Only SNPs from “Adafuel” linkage map were selected as they covered the eight-linkage group. Conversely, only four groups were constructed for “Flordaguard”. Currently this is the only dataset in PrunusMap featuring genetic positions.

For almond, the Axiom 60K SNP array was developed from whole-genome resequencing data of 81 almond genomes (Duval et al., 2023).

#### 2.2.4 Annotations

Gene datasets were enriched with functional annotations from InterPro and Pfam databases (Mistry et al., 2021; Paysan-Lafosse et al., 2023). To ensure the highest accuracy, annotations were limited to references *Prunus persica* (Pp_JGI_V2) and *Prunus dulcis* (Pd_Texas_GDR_V2).

### 2.3 PrunusMap Commands: Navigating the Toolkit

PrunusMap offers a variety of Web and standalone tools, which are summarized below. While the former are publicly accessible, standalone tools require local installation and configuration of PrunusMap. Check the repository https://github.com/eead-csic-compbio/prunusmap_web and the help section at https://prunusmap.eead.csic.es/prunusmap/help for more details.

#### 2.3.1 Web-based tools

Markers can be searched using different input: FASTA sequences, IDs or positions. This is facilitated by the following standalone tools, which can examine up to 10,000 entries on single or multiple maps.

- bmap_align”: aligns FASTA-formatted sequences to reference databases using GMAP, BLASTN, or both (Sayers et al., 2022; Wu & Watanabe, 2005). Initially, queries are searched using GMAP. If no matches are found, BLASTN takes over. This iteration continues until either all queries are aligned, or no additional databases are available. The default parameters are set as minimum identity=98% and minimum coverage=95% but can be customized to suit user’s requirements.
- bmap_find”: takes a list of query IDs and retrieves their alignment positions from the pre-computed datasets listed in **Table 1**.
- bmap_locate”: locates features (genes and/or markers) based on their position within chromosomes or scaffolds.

#### 2.3.2 Standalone version

Secondary tools such as “bmap_align_to_db” and “bmap_align_to_map” are only available in the standalone version which can be installed following the instructions in the GitHub repository. They provide in-depth alignment results which are not reported on the Web.

#### 2.3.3 PrunusMap output

The output is provided through the Web interface or conveniently sent to an email address for ease of interpretation and storage. Within the Web interface, the results are displayed in two formats: as a graphical representation of the genome, emphasizing the query locations and as a downloadable CSV file (**Figure 1**). PrunusMap also provides additional tables showcasing the positions of nearby markers, genes, or proteins.

The search radius for neighboring features can be tailored and defined either in base pairs (bp) or in centimorgans (cM), depending on the underlying selected map (**Figure S2**). Additional details are provided in the help section https://prunusmap.eead.csic.es/prunusmap/help.

#### 2.3.4 Showcase analysis

To benchmark PrunusMap, we analyzed relevant markers documented in the literature. For instance, (Fleming et al., 2022) reported two quantitative trait loci (QTLs) linked to fruit resistance to bacterial spot infection (“Xap.PpOC-1.2” and “Xap.PpOC-1.6”), sitting on chromosomes 1 and 6. Eleven SNPs in close proximity to these QTLs were used to design KASP markers, resulting in 44% reduction in seedling planting (Fleming et al., 2022). In this context, we tested “Find markers” Web tool in order to locate the eleven SNPs that correspond to following 9K peach array markers: SNP_IGA_39717, SNP_IGA_40295, SNP_IGA_43384, SNP_IGA_46754, SNP_IGA_680615, SNP_IGA_680882, SNP_IGA_680909, SNP_IGA_680953, SNP_IGA_681081, SNP_IGA_681113 and SNP_IGA_681119.

## 3. Results

### 3.1 Comparing GMAP and BLASTN

The performance of GMAP and BLASTN was assessed by aligning marker sets against different versions of the peach and almond reference genomes (see **Figure 3**). Note that the peach references from NCBI and Phytozome are identical but have distinct annotations. For “Adafuel” and 9K datasets both aligners achieved similar accuracy, mapping nearly the same number of sequences. Specifically, GMAP retrieved only 2 and 31 additional positions, respectively. Regarding the 16K array, with a total of 16,026 markers (gray bar **in Figure 3.A**), GMAP mapped 15,834 sequences, compared to 14,337 from BLASTN.

**Figure 3.**
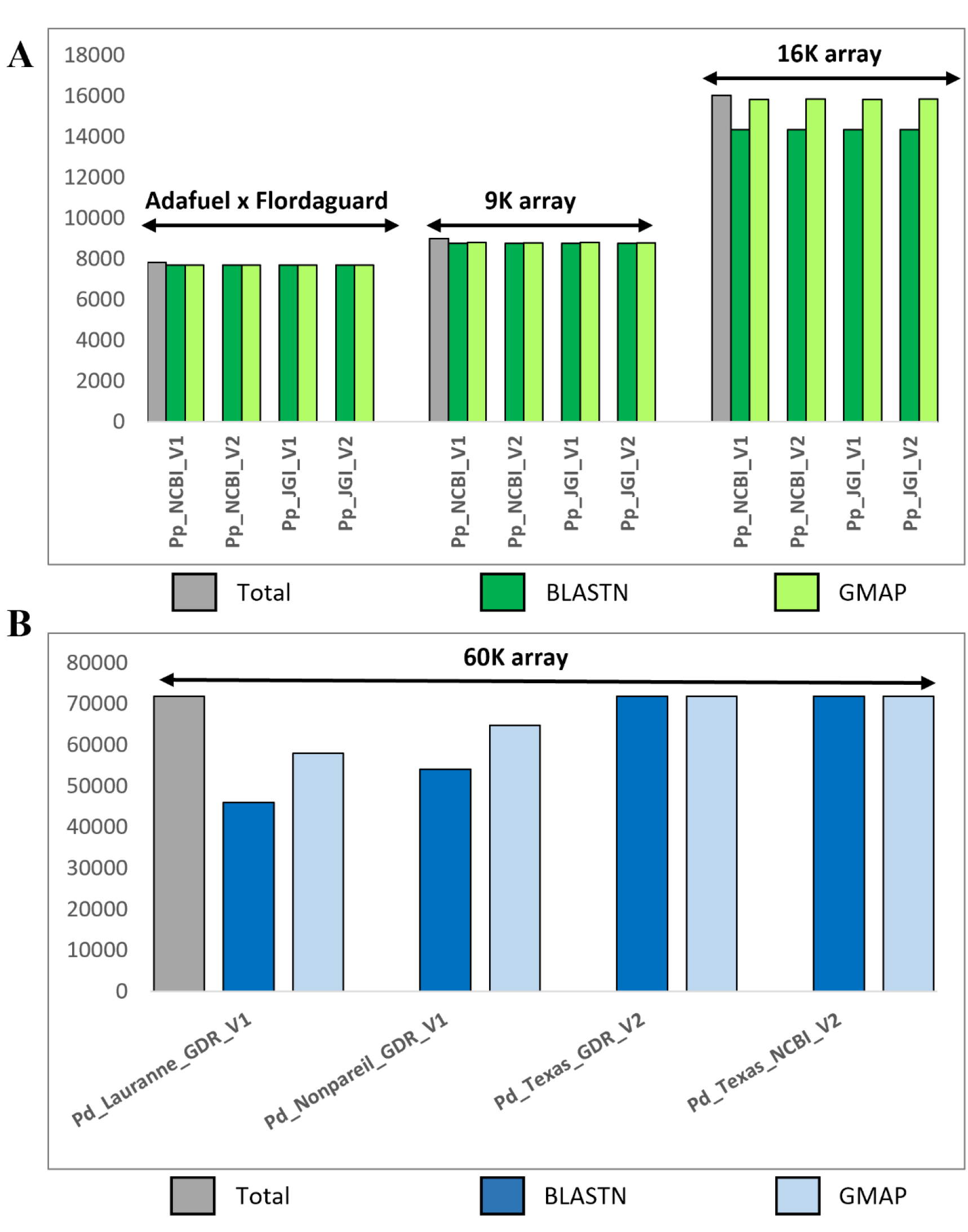
Performance comparison of GMAP and BLASTN aligners using genetic markers and different versions of *Prunus persica* **(A)** and *Prunus dulcis* **(B)** genome references. The x-axis refers to the different databases (maps according to PrunusMap terminology) used for the alignment while the y-axis corresponds to the number of sequences. Gray bars correspond to the total number of sequence markers to be aligned, dark green and dark blue bars correspond to the aligned hits using BLASTN. Light green and light blues bars correspond to aligned hits with GMAP.

For *Prunus dulcis*, a total of 71,843 markers from the 60K Axiom array were aligned to the Texas, Lauranne and Nonpareil databases. As shown in **Figure 3.B**, most markers matched the Texas database, regardless of the aligner. This is probably due to the array design, which involved the alignment of re-sequenced raw reads against Texas prior to SNP calling (Duval et al., 2023). Overall, GMAP demonstrated superior performance by successfully aligning 71,841, 57,833, and 64,731 markers compared to BLASTN, which aligned (71,838, 55,984, and 54,016) to the Texas, Lauranne, and Nonpareil databases, respectively. Additionally, GMAP exhibited faster processing times (seconds vs minutes) compared to BLASTN. Unmapped queries can be attributed to the stringent sequence identity and query coverage thresholds, set by default at 98% and 95%, respectively. Hits below these cutoffs are deemed unreliable and are discarded from the datasets.

To further evaluate alignment performance, we examined transcripts from the peach reference transcriptome (RefTrans_V1; 23,390). These sequences were downloaded from the GDR and derived from publicly available RNA-Seq and EST data sets. **Figure 4** shows that BLAST matched 4,902 and 4,952 transcripts, compared to 20,482 and 20,502 matched by GMAP.

**Figure 4.**
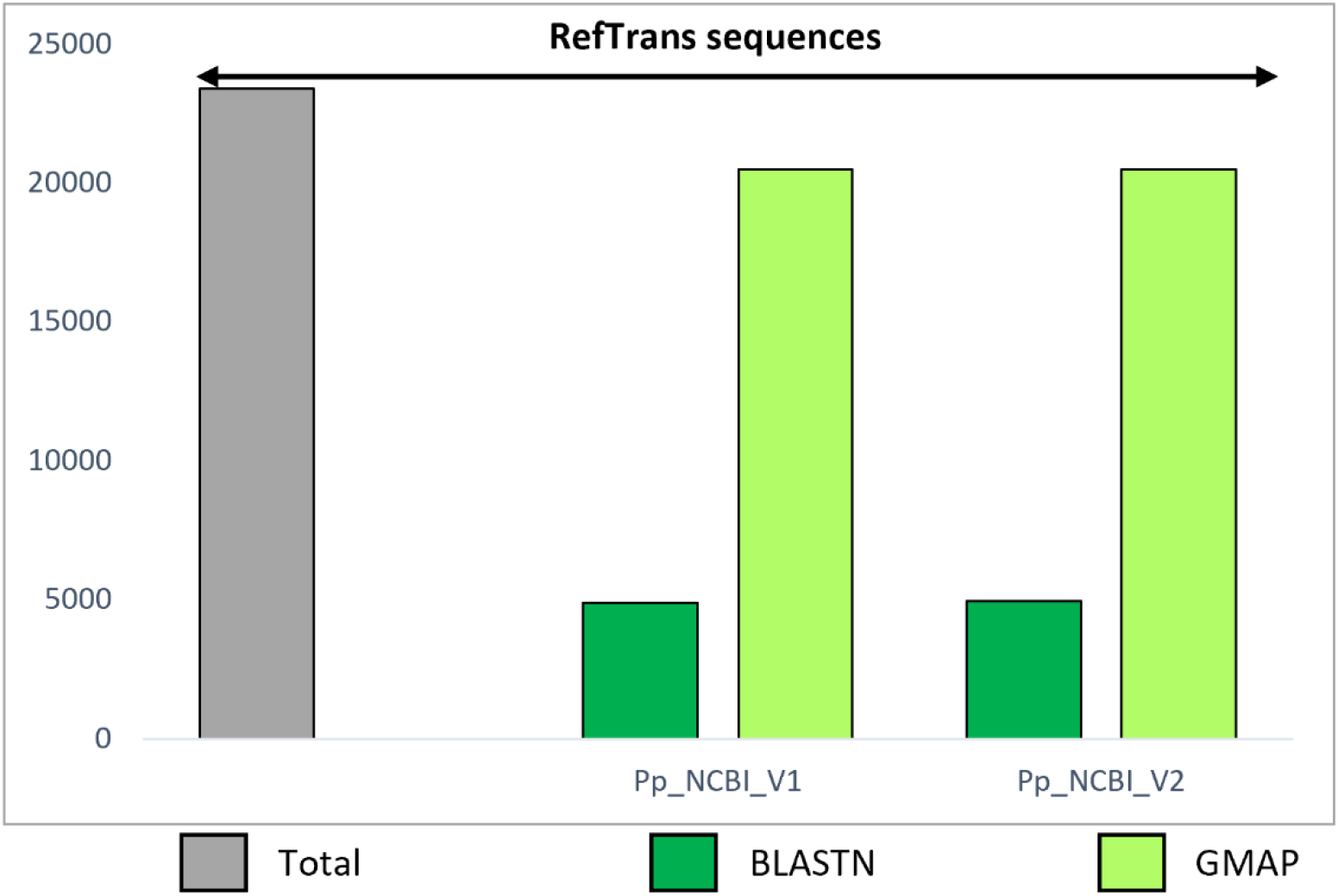
Performance comparison of GMAP and BLASTN aligners using long transcripts and *Prunus persica* genome assemblies. The x-axis refers to the peach genome versions (maps according to PrunusMap terminology), while the y-axis corresponds to the total number of transcripts to be aligned.

### 3.2 Benchmarking the accuracy of PrunusMap

“Adafuel” marker sequences were aligned against the different databases of *Prunus persica* using “bmap_align”. The resulting physician positions (in Mbp) were plotted against their genetic positions (in cM), as reported by (Guajardo et al., 2022). Notably, high collinearity was evident across all chromosomes, with *Pearson* correlation coefficients ≥ 0.96 for reference JGI_V2 (**Figure 5.A**). Overlapping gaps between genetic and physical positions were mainly observed in chromosomes Pp02, Pp07 and Pp08. These are explained by the uneven distribution of markers mapped in the “Adafuel × Flordaguard” population (Guajardo et al., 2022). Note that there are no markers mapping on the short arm of Pp04. Interestingly, when JGI_V1 was used as reference, inversions in scaffolds 1, 6 and 7 were revealed, as confirmed by (Verde et al., 2017) (**Figure 5.B**). Similarly, a misplaced contig appears near the centromere of scaffold 4. According to our results, these assembly conformations were corrected in V2, and 60 chromosomal SNPs previously located on unplaced scaffolds were also rectified. Comparable results were obtained for NCBI references, plotted in **Figures S3** and **S4**.

**Figure 5.**
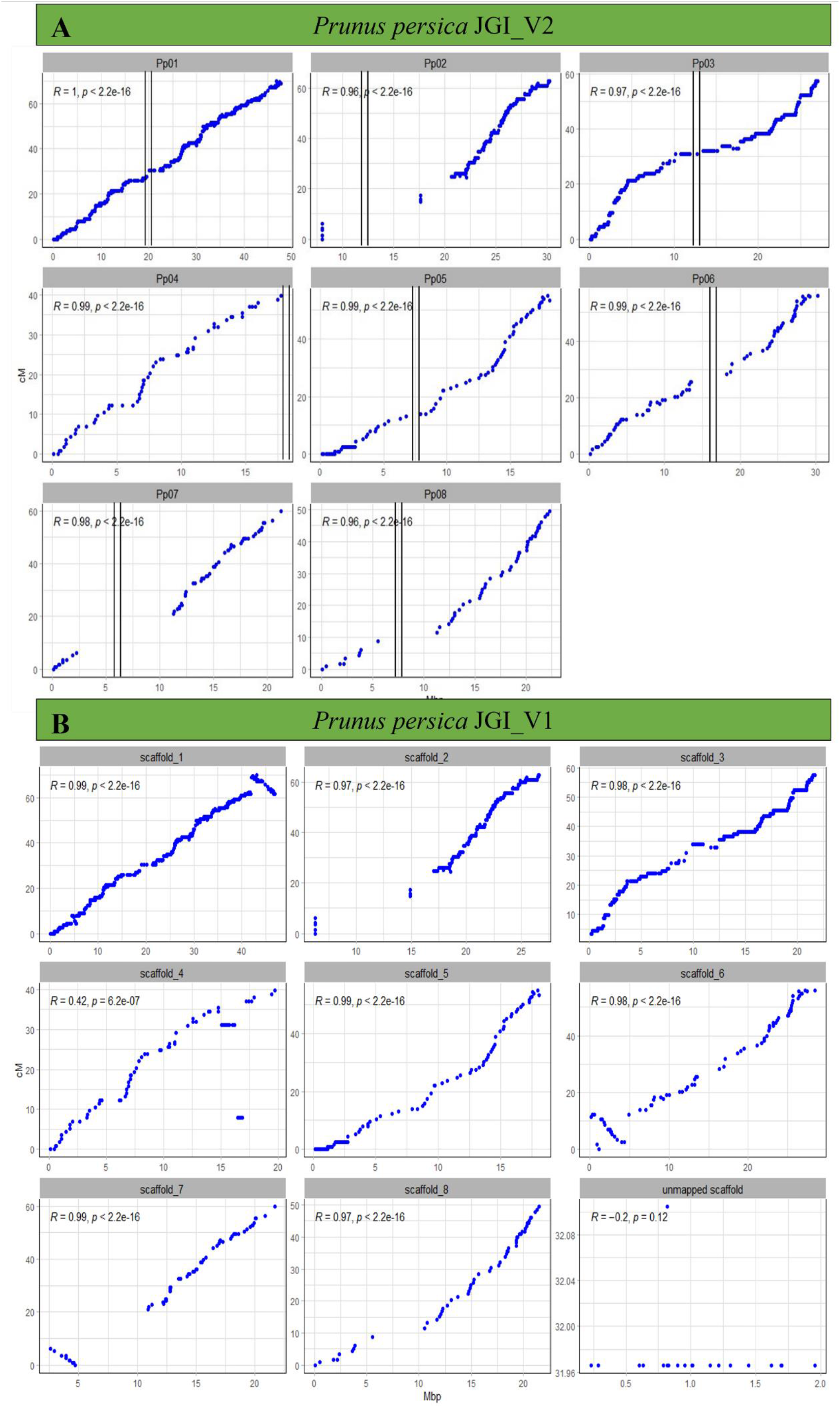
Relationship between genetic and physical position of “Adafuel” SNP markers within each pseudomolecule (chromosomes). For *Prunus persica* JGI_V2 **(A)**, pseudomolecules are referred to as Pp01 to Pp08, while in *Prunus persica* JGI_V1 **(B)**, they are labeled as scaffold_1 to scaffold_2. SNP markers were plotted according to their physical position on peach genome reference (x-axis), and their genetic position retrieved from “Adafuel” linkage map (y-axis). Vertical bars indicate putative position of the centromeres and R values correspond to the *Pearson* correlation.

### 3.3 Examining the insights provided by PrunusMap

“Find markers” was used to assess a list of 11 relevant markers associated with bacterial spot resistance in peach. As depicted in **Figure 6**, 10 out 11 were successfully mapped to their corresponding chromosomes (Pp01 and Pp06 according to the JGI_V2 reference genome). SNP_IGA_46754 remained unmapped for not meeting the default alignment quality cutoffs. The screenshot in the **Figure 6** includes the physical positions of mapped markers, neighboring genes and their functional annotations. Several genes encode disease resistance proteins with LRR and NB-ARC domains (*Prupe*.*1G165300, Prupe*.*6G243700* and *Prupe*.*6G243800*), known as major disease resistance genes in plants. For instance, in peach they have been reported to be involved in pathogen recognition and innate immune responses (Fu et al., 2021).

**Figure 6.**
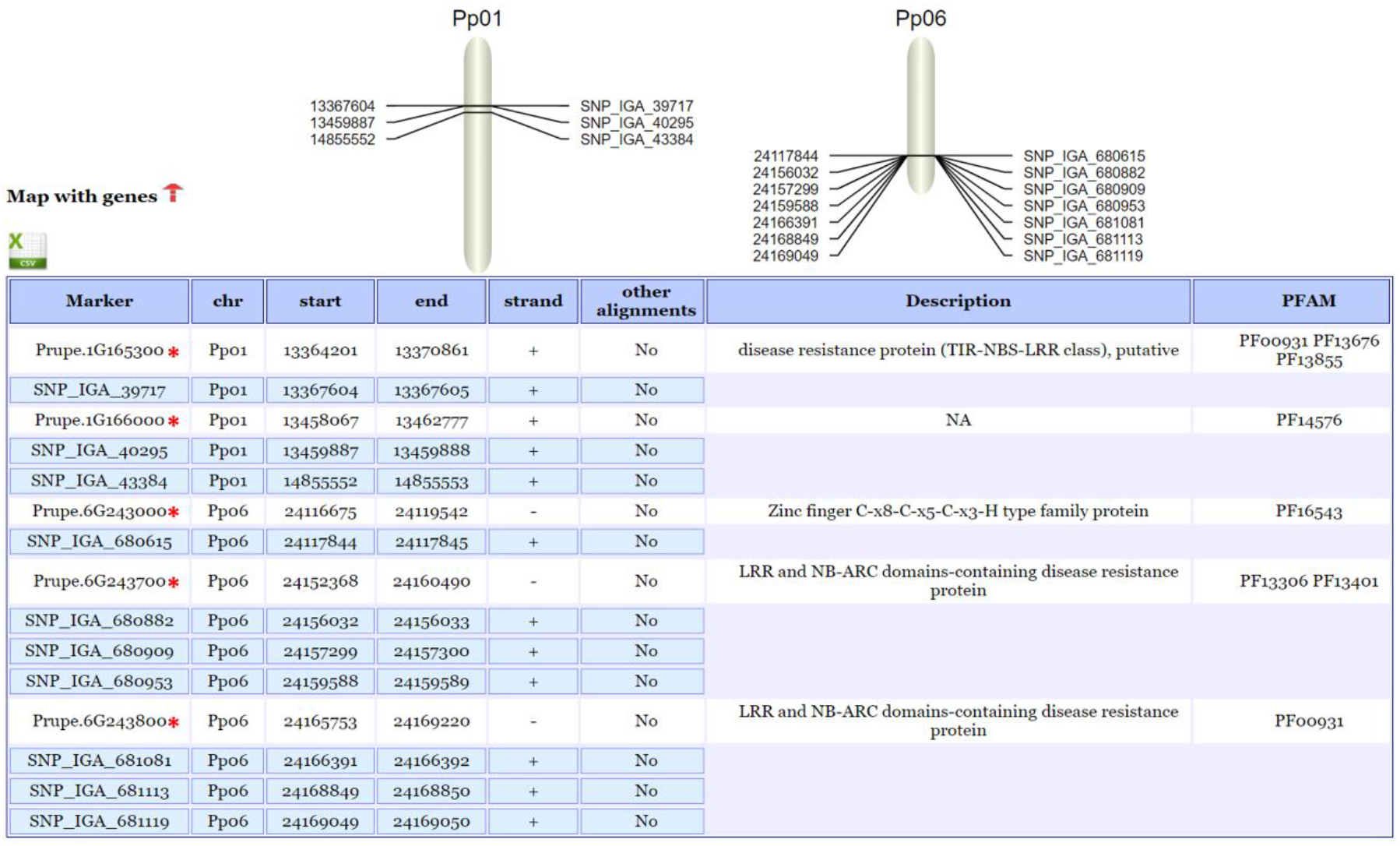
Illustration of PrunusMap functionalities. Genes marked with red asterisk correspond to nearby genes identified through the “Find marker” tool.

## 4. Discussion

In the following sections we compare PrunusMap and other tools offering similar functionalities.

### 4.1 GDR and PrunusMap

GDR serves as a central hub of the Rosaceae family. As a public repository, it provides access to multiple versions of genome assemblies enriched with gene descriptions, InterPro domains, GO and KEGG pathways terms (Jung et al., 2019). Moreover, it hosts collections of expressed sequence tags (ESTs), full-length transcripts, metabolic pathways, maps and quantitative and mendelian trait loci linked to agronomically significant traits (Jung et al., 2019). Beyond its role as a repository, GDR provides analytical tools to explore genetic and genomic data (search, sequence retrieval, BLAST, synteny viewer, map viewer, BIMS).

While sharing similar purposes, PrunusMap is specifically designed to facilitate the identification of *Prunus* markers on both physical and genetic maps, providing a streamlined analysis that includes a detailed list of nearby genes and proteins. Although this functionality may seem to be fulfilled by GDR, there are key differences incorporated in PrunusMap.

First, regarding sequence homology searches, GDR relies exclusively on BLAST, whereas PrunusMap employs the GMAP as its default aligner complemented with BLASTN. Based on our findings, GMAP consistently outperformed BLASTN in locating marker sequences (**Figure 3**). This performance gain is further accentuated when mapping transcripts, which require intron-aware alignments. These results agree with those reported in barley (Cantalapiedra et al., 2015).

Second, when searching for molecular markers, GDR supports an extensive list of parameters such as marker type, name, array, organism, chromosome, map, trait, and citation. PrunusMap further refined its search capabilities to align with our vision, offering graphical visualization of marker locations on their corresponding chromosomes. This view is accompanied by a table summarizing the start and end positions of the markers, the strand, and crucially, a list of annotated genes and proteins residing nearby. The strand orientation, absent in GDR, is particularly valuable for breeders as it can assist in designing PCR primers or KASP markers, predicting missense mutations and defining haplotypes. Finally, by displaying nearby genes on the same results table, PrunusMap eliminates the need to conduct multiple searches and navigate from one webpage to another. This streamlined approach enhances efficiency and simplifies the process for researchers and breeders alike.

Third, a significant advantage of PrunusMap is the ability to map the same input queries across different genome versions of the same species (*P. persica*), as well as on different cultivars (*P. dulcis*). This capability way PrunusMap helps address the gene nomenclature discrepancy that already exists between different genome versions, such as those in the NCBI and JGI. Additionally, many published works often tend to focus on a single cultivar, resulting in other cultivars being overlooked. For instance, a marker closely linked to a desirable trait in one cultivar may exhibit a different position or allele in another cultivar. PrunusMap can handle this imbalance enabling breeders to explore several references in the same search. This, in turn facilitates the identification of new sources of desirable traits. For example, the ability to map the 60K Axiom markers on three different almond references (Texas, Lauranne and Nonpareil) would enable breeders to easily explore the genetic diversity and relationships among these three cultivars.

Fourth, PrunusMap streamlines retrieving the physical and genetic positions of all markers from a given genetic map in a single operation, as shown here with “Adafuel” markers, which supports the analysis of genetic maps in a single step.

Finally, PrunusMap further enhances user experience by enabling the option to send search results via email. This feature is particularly helpful for large scale analyses that may require extended processing times.

Overall, both GDR and PrunusMap serve as valuable tools, each offering distinct, yet complementary features tailored to different user preferences.

### 4.2. PeachVar-DB and PrunusMap

PeachVar-DB is a valuable resource for exploring the genetic makeup of a collection comprising 121 peach accessions and 21 wild relatives from the *Amygdalus* subgenus derived from the re-sequencing (Cirilli et al., 2018). Users can get a broad overview by selecting a specific accession, delve into a specific genome region, or conduct a targeted gene-level analysis by providing the gene ID and features such as 5’UTR, 3’UTR, CDS, or primary transcript. However, unlike PrunusMap, when selecting an accession or chromosome region, PeachVar-DB does not display information on nearby genes alongside the genetic variants. As with GDR, PeachVar-DB utilizes BLASTN for sequence similarity comparison; however, it does not feature a graphical visualization of the mapping results. Furthermore, in contrast to PrunusMap, PeachVar-DB does not support multi-query searches and file upload functionalities. Finally, it exclusively presents information on markers aligned to the peach reference genome v2.0.

## 5. Conclusions and future directions

PrunusMap was developed to empower *Prunus* researchers with user-friendly analysis tools to support decision-making and accelerate breeding goals. We anticipate it will serve as a valuable tool for breeders in combination with GDR and PeachVar-DB. Furthermore, we expect PrunusMap to be continuously updated and expanded to cover other *Prunus* species upon demand from users. We welcome feedback and suggestions at.

## Funding and Acknowledgments

This work was funded by CSIC [grants 2020AEP119 & FAS2022_052], the Spanish Research Agency [grants AGL2017-83358-R MCIN/AEI/10.13039/501100011033 & “A way of making Europe”] and the Government of Aragón [grants A09_23R, A10_23R & PhD contract to N. Ksouri 2018-2023], which were co-financed with FEDER funds. The authors would like to thank Dr V. Guajardo for providing data of “Adafuel” SNPs.

## Contributions

Y.G. and B.C-M. conceived the project and its components. M-A.M. provided genetic data, B.C-M. and N.K. configured the application and maintain the server, N.K. created the datasets and wrote the draft manuscript, B.C-M. and Y.G. discussed and revised the manuscript. All authors read and approved the final manuscript.

## Data Availability

PrunusMap is freely accessible at https://prunusmap.eead.csic.es. Instructions for configuring and using the Web application can be found at https://github.com/eead-csic-compbio/prunusmap_web

## References

Alioto, T., Alexiou, K. G., Bardil, A., Barteri, F., Castanera, R., Cruz, F., Dhingra, A., Duval, H., Fernández i Martí, Á., Frias, L., Galán, B., García, J. L., Howad, W., Gómez-Garrido, J., Gut, M., Julca, I., Morata, J., Puigdomènech, P., Ribeca, P., & Arús, P. (2020). Transposons played a major role in the diversification between the closely related almond and peach genomes: results from the almond genome sequence. Plant Journal, 101(2), 455–472. 10.1111/tpj.14538

Aranzana, M. J., Decroocq, V., Dirlewanger, E., Eduardo, I., Gao, Z. S., Gasic, K., Iezzoni, A., Jung, S., Peace, C., Prieto, H., Tao, R., Verde, I., Abbott, A. G., & Arús, P. (2019). Prunus genetics and applications after de novo genome sequencing: achievements and prospects. In Horticulture Research (Vol. 6, Issue 58, pp. 1–25). Nature Publishing Group. 10.1038/s41438-019-0140-8

Araújo, M. de C., Rodrigues, P., Filho, S. A., & Clement, C. R. (2010). Genetic variability in the peach palm genebank examined with RAPD markers. Crop Breeding and Applied Biotechnology, 10, 211–217.

Boratyn, G. M., Camacho, C., Cooper, P. S., Coulouris, G., Fong, A., Ma, N., Madden, T. L., Matten, W. T., McGinnis, S. D., Merezhuk, Y., Raytselis, Y., Sayers, E. W., Tao, T., Ye, J., & Zaretskaya, I. (2013). BLAST: a more efficient report with usability improvements. Nucleic Acids Research, 41, 29–33. 10.1093/nar/gkt282

Butiuc-Keul, A., Coste, A., Postolache, D., Laslo, V., Halmagyi, A., Cristea, V., & Farkas, A. (2022). Molecular characterization of Prunus cultivars from Romania by microsatellite markers. Horticulturae, 8(4). 10.3390/horticulturae8040291

Cantalapiedra, C. P., Boudiar, R., Casas, A. M., Igartua, E., & Contreras-Moreira, B. (2015). BARLEYMAP: physical and genetic mapping of nucleotide sequences and annotation of surrounding loci in barley. Molecular Breeding, 35(13), 1–13. 10.1007/s11032-015-0253-1

Cirilli, M., Flati, T., Gioiosa, S., Tagliaferri, I., Ciacciulli, A., Gao, Z., Gattolin, S., Geuna, F., Maggi, F., Bottoni, P., Rossini, L., Bassi, D., Castrignanò, T., & Chillemi, G. (2018). PeachVar-DB: A curated collection of genetic variations for the interactive analysis of peach genome data. Plant & Cell Physiology, 59(1), 1–9. 10.1093/pcp/pcx183

Cirilli, M., Rossini, L., Geuna, F., Palmisano, F., Minafra, A., Castrignanò, T., Gattolin, S., Ciacciulli, A., Babini, A. R., Liverani, A., & Bassi, D. (2017). Genetic dissection of Sharka disease tolerance in peach (P. persica L. Batsch). BMC Plant Biology, 17(1), 1–15. 10.1186/s12870-017-1117-0

D’Amico-Willman, K. M., Ouma, W. Z., Meulia, T., Sideli, G. M., Gradziel, T. M., & Fresnedo-Ramirez, J. (2022). Whole-genome sequence and methylome profiling of the almond [Prunus dulcis (Mill.) D.A. Webb] cultivar ‘Nonpareil.’ G3: Genes, Genomes, Genetics, 12(5), 1–9. 10.1093/g3journal/jkac065

Duval, H., Coindre, E., Ramos-Onsins, S. E., Alexiou, K. G., Rubio-Cabetas, M. J., Martínez-García, P. J., Wirthensohn, M., Dhingra, A., Samarina, A., & Arús, P. (2023). Development and evaluation of an AxiomTM 60K SNP array for almond (Prunus dulcis). Plants, 12(242), 1–12. 10.3390/plants12020242

Fleming, M. B., Miller, T., Fu, W., Li, Z., Gasic, K., & Saski, C. (2022). Ppe.XapF: High throughput KASP assays to identify fruit response to Xanthomonas arboricola pv. pruni (Xap) in peach. PLoS ONE, 17, 1–17. 10.1371/journal.pone.0264543

Fu, W., da Silva Linge, C., & Gasic, K. (2021). Genome-wide association study of Brown Rot (Monilinia spp.) tolerance in peach. Frontiers in Plant Science, 12, 1–14. 10.3389/fpls.2021.635914

Gasic et al. 2022. Unpublished. IRSC 16K markers downloaded from the GDR.

Gillen, A. M., & Bliss, F. A. (2005). Identification and mapping of markers linked to the Mi gene for root-knot nematode resistance in peach. Journal of the American Society for Horticultural Science, 130(1), 24–33.

Goodstein, D. M., Shu, S., Howson, R., Neupane, R., Hayes, R. D., Fazo, J., Mitros, T., Dirks, W., Hellsten, U., Putnam, N., & Rokhsar, D. S. (2012). Phytozome: A comparative platform for green plant genomics. Nucleic Acids Research, 40, D1178–D1186. 10.1093/nar/gkr944

Guajardo, V., Martínez-García, P. J., Solís, S., Calleja-Satrustegui, A., Saski, C., & Moreno, M. Á. (2022). QTLs identification for iron chlorosis in a segregating peach– almond progeny through double-digest sequence-based genotyping (SBG). Frontiers in Plant Science, 13(872208), 1–14. 10.3389/fpls.2022.872208

Hellegouarch, S. (2007). CherryPy: Essentials rapid Python Web application development design, develop, test, and deploy your Python Web applications easily. Packt publishing Ltd. http://www.visionwt.com

Jung, S., Jesudurai, C., Staton, M., Du, Z., Ficklin, S., Cho, I., Abbott, A., Tomkins, J., & Main, D. (2004). GDR (Genome Database for Rosaceae): Integrated web resources for Rosaceae genomics and genetics research. BMC Bioinformatics, 5(130), 1–8. 10.1186/1471-2105-5-130

Jung, S., Lee, T., Cheng, C.-H., Buble, K., Zheng, P., Yu, J., Humann, J., Ficklin, S. P., Gasic, K., Scott, K., Frank, M., Ru, S., Hough, H., Evans, K., Peace, C., Olmstead, M., DeVetter, L. W., McFerson, J., Coe, M., … Main, D. (2019). 15 years of GDR: New data and functionality in the genome database for Rosaceae. Nucleic Acids Research, 47, D1137–D1145. 10.1093/nar/gky1000

Jung, S., Staton, M., Lee, T., Blenda, A., Svancara, R., Abbott, A., & Main, D. (2008). GDR (Genome Database for Rosaceae): Integrated web-database for Rosaceae genomics and genetics data. Nucleic Acids Research, 36, 1034–1040. 10.1093/nar/gkm803

Li, H. (2023). Protein-to-genome alignment with miniprot. Bioinformatics, 39(1), 1–6. 10.1093/bioinformatics/btad014

Mistry, J., Chuguransky, S., Williams, L., Qureshi, M., Salazar, G. A., Sonnhammer, E. L. L., Tosatto, S. C. E., Paladin, L., Raj, S., Richardson, L. J., Finn, R. D., & Bateman, A. (2021). Pfam: The protein families database in 2021. Nucleic Acids Research, 49, D412–D419. 10.1093/nar/gkaa913

Paysan-Lafosse, T., Blum, M., Chuguransky, S., Grego, T., Pinto, B. L., Salazar, G. A., Bileschi, M. L., Bork, P., Bridge, A., Colwell, L., Gough, J., Haft, D. H., Letunić, I., Marchler-Bauer, A., Mi, H., Natale, D. A., Orengo, C. A., Pandurangan, A. P., Rivoire, C., & Bateman, A. (2023). InterPro in 2022. Nucleic Acids Research, 51, D418–D427. 10.1093/nar/gkac993

Sánchez-Pérez, R., Pavan, S., Mazzeo, R., Moldovan, C., Cigliano, R. A., Cueto, J. Del, Ricciardi, F., Lotti, C., Ricciardi, L., Dicenta, F., López-Marqués, R. L., & Lindberg Møller, B. (2019). Mutation of a bHLH transcription factor allowed almond domestication. Science, 364, 1095–1098. 10.1126/science.aav8197

Sayers, E. W., Bolton, E. E., Brister, J. R., Canese, K., Chan, J., Comeau, D. C., Connor, R., Funk, K., Kelly, C., Kim, S., Madej, T., Marchler-Bauer, A., Lanczycki, C., Lathrop, S., Lu, Z., Thibaud-Nissen, F., Murphy, T., Phan, L., Skripchenko, Y., & Sherry, S. T. (2022). Database resources of the national center for biotechnology information. Nucleic Acids Research, 50, D20–D26. 10.1093/nar/gkab1112

Verde, I., Abbott, A. G., Scalabrin, S., Jung, S., Shu, S., Marroni, F., Zhebentyayeva, T., Dettori, M. T., Grimwood, J., Cattonaro, F., Zuccolo, A., Rossini, L., Jenkins, J., Vendramin, E., Meisel, L. A., Decroocq, V., Sosinski, B., Prochnik, S., Mitros, T., & Rokhsar, D. S. (2013). The high-quality draft genome of peach (Prunus persica) identifies unique patterns of genetic diversity, domestication and genome evolution. Nature Genetics, 45(5), 487–494. 10.1038/ng.2586

Verde, I., Bassil, N., Scalabrin, S., Gilmore, B., Lawley, C. T., Gasic, K., Micheletti, D., Rosyara, U. R., Cattonaro, F., Vendramin, E., Main, D., Aramini, V., Blas, A. L., Mockler, T. C., Bryant, D. W., Wilhelm, L., Troggio, M., Sosinski, B., Aranzana, M. J., & Peace, C. (2012). Development and evaluation of a 9K SNP array for peach by internationally coordinated SNP detection and validation in breeding germplasm. PLoS ONE, 7(4), 1–13. 10.1371/journal.pone.0035668

Verde, I., Jenkins, J., Dondini, L., Micali, S., Pagliarani, G., Vendramin, E., Paris, R., Aramini, V., Gazza, L., Rossini, L., Bassi, D., Troggio, M., Shu, S., Grimwood, J., Tartarini, S., Dettori, M. T., & Schmutz, J. (2017). The Peach v2.0 release: high-resolution linkage mapping and deep resequencing improve chromosome-scale assembly and contiguity. BMC Genomics, 18(225), 1–18. doi:10.1186/s12864-017-3606-9

Viruel, M. A., Messeguer, R., de Vicente J Garcia-Mas, M. C., Puigdom, P., Vargas P Arfis, nech F., Garcia-Mas, J., & Vargas, F. (1995). A linkage map with RFLP and isozyme markers for almond. Theoretical and Applied Genetics, 964–971. 10.1007/BF00223907

Wang, Y., Georgi, L. L., Reighard, G. L., Scorza, R., & Abbott, A. G. (2002). Genetic mapping of the evergrowing gene in peach [Prunus persica (L.) Batsch]. Journal of Heredity, 93(5), 352–358. https://academic.oup.com/jhered/article/93/5/352/2187321

Wu, T. D., & Watanabe, C. K. (2005). GMAP: A genomic mapping and alignment program for mRNA and EST sequences. Bioinformatics, 21(9), 1859–1875. 10.1093/bioinformatics/bti310

