## Supplementary Figures for "Mapping the genomic landscape of peach and almond with PrunusMap"

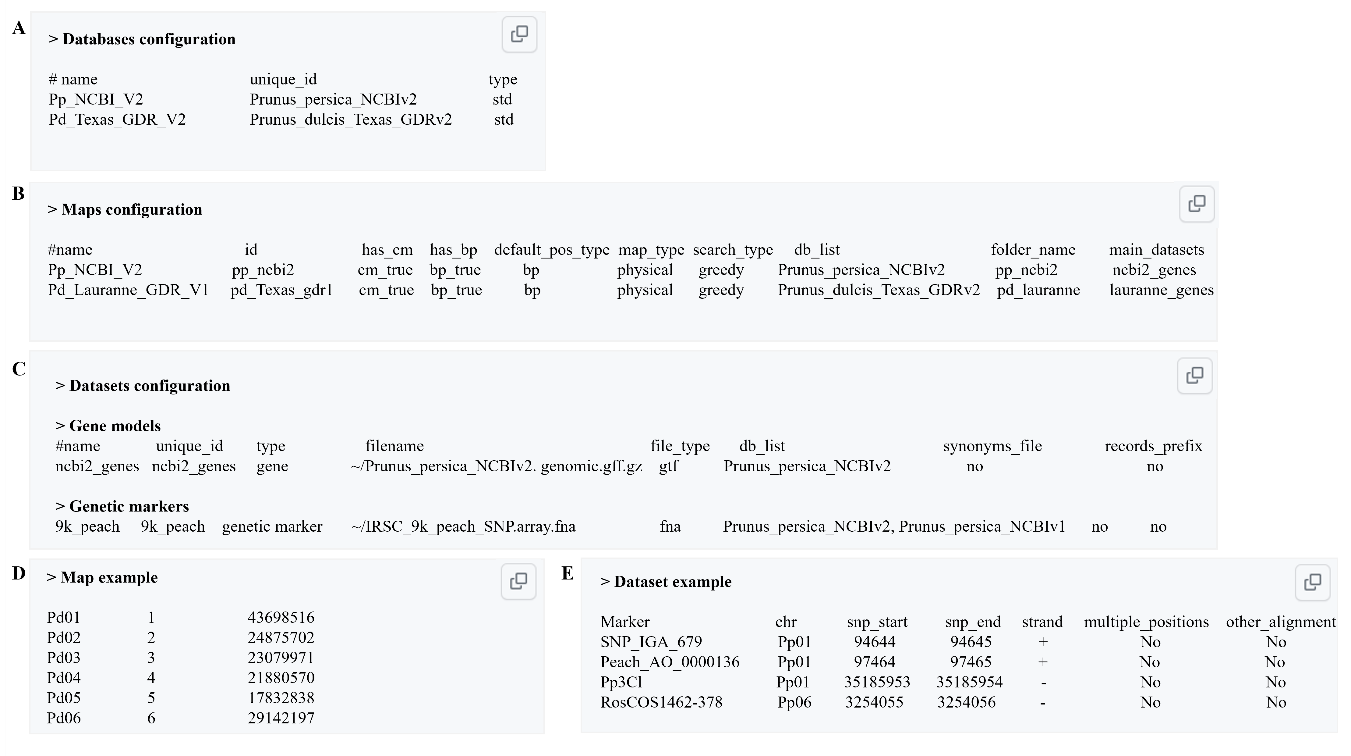

**Supplemental Figure S1**. An overview of biological resource configuration in the PrunusMap framework. Panels **A**, **B** and **C** represent the file structure of databases, maps and datasets respectively. Panels **D** and **E** showcase examples of maps and datasets.

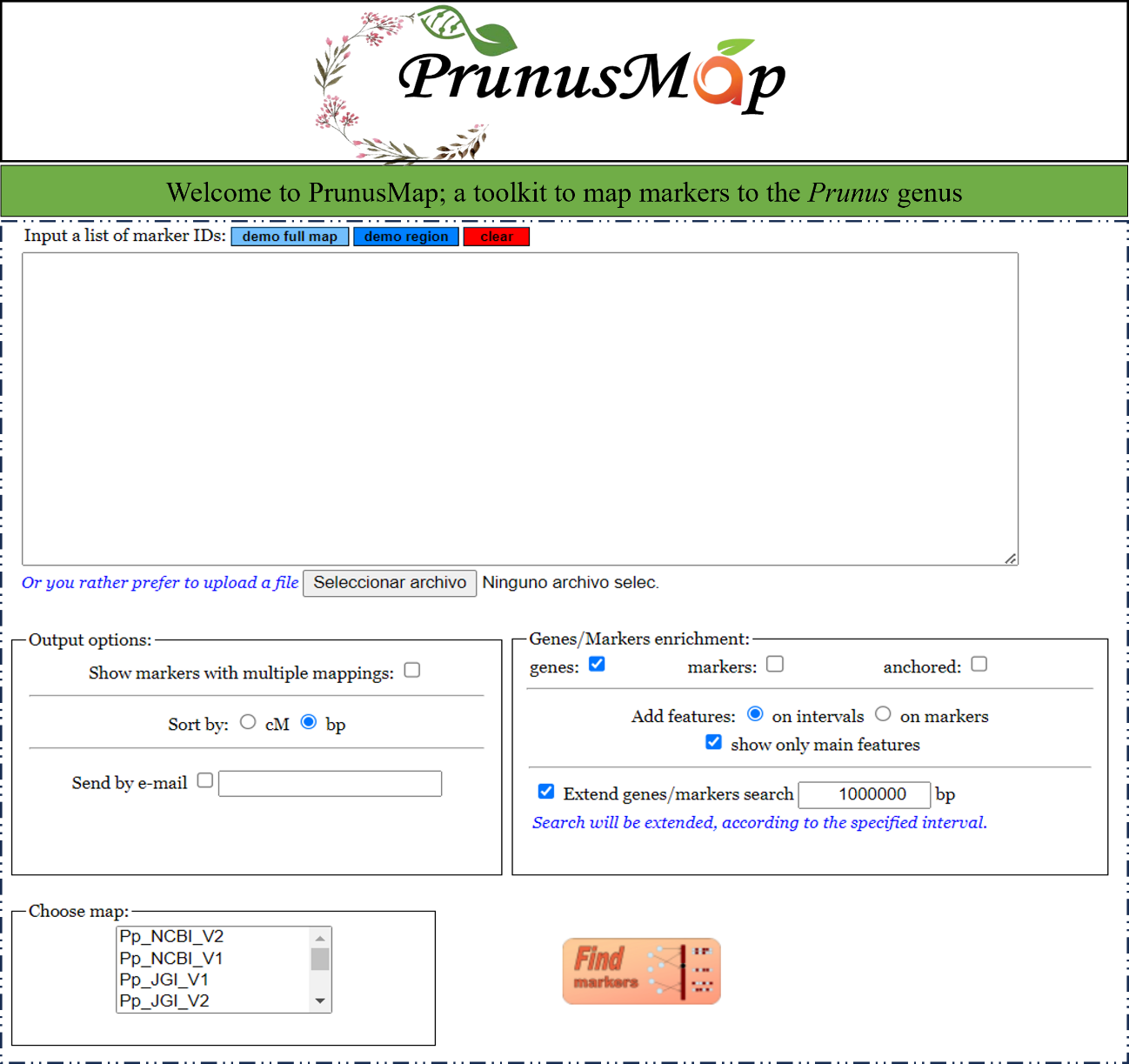

**Supplemental Figure S2**. Screen capture of PrunusMap Web portal. Queries can be entered directly in the input box or uploaded as a file. The output options box gives the users the option to provide their email and to sort the results by their physical (bp) or genetic positions (cM). The Genes/Marker enrichment panel allows users to retrieve the adjacent features (genes, markers and/or proteins) within a user-customized interval.

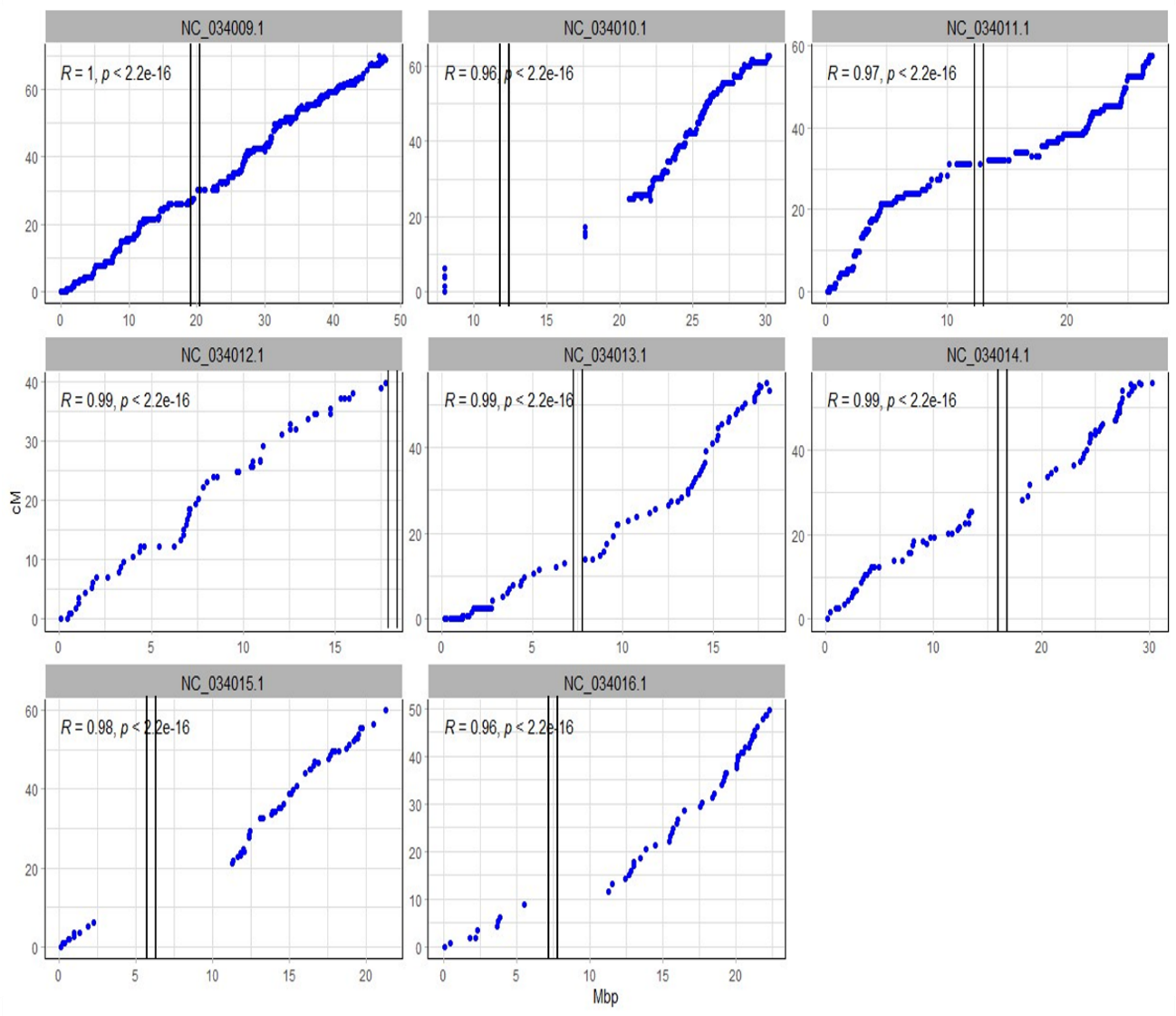

**Supplemental Figure S3.** Relationship between genetic and physical position of “Adafuel” SNP markers mapped to peach reference genome NCBI_V2. Pseudomolecules are referred to as NC_034009.1 to NC_034016.1. Markers were plotted according to their physical position in Mbp on peach NCBI_V2 (x-axis) and their genetic position in cM (y-axis). Vertical bars indicate putative position of the centromeres.

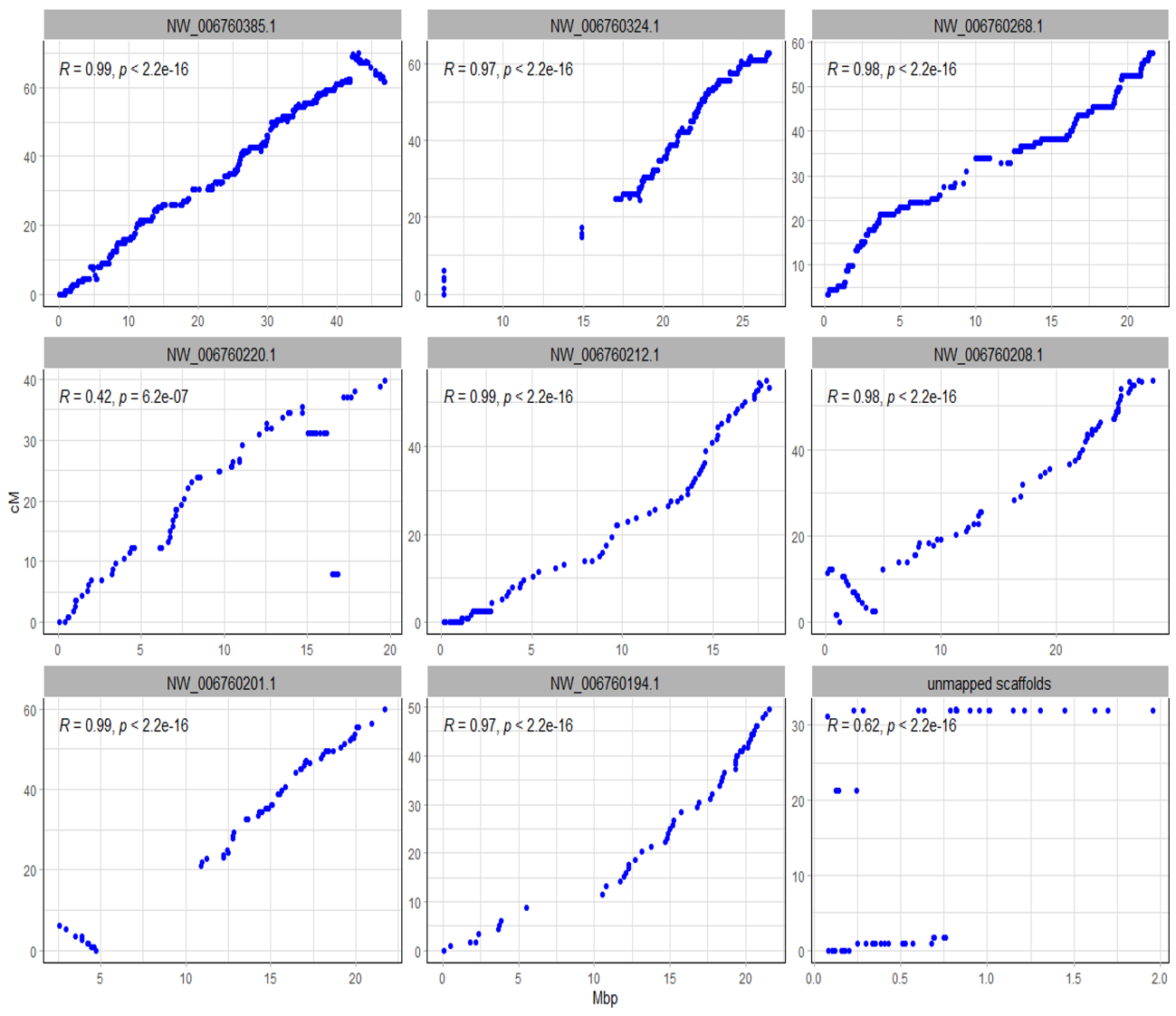

**Supplemental Figure S4.** Relationship between genetic and physical position of “Adafuel” SNP markers mapped to peach reference genome NCBI_V1. Scaffolds are referred to as NW_006760385.1 and so forth. Markers were plotted according to their physical position in Mbp on peach NCBI_V1 (x-axis) and their genetic position in cM (y-axis) from the “Adafuel” genetic map.
